# A TORC1-PHR1 signaling axis regulates phosphorus starvation and immunity signaling network in Arabidopsis

**DOI:** 10.1101/2024.11.18.624151

**Authors:** Prakhar Awasthi, K Muhammed Jamsheer, Manoj Kumar, Shruti Mishra, Archna Tiwari, Sunita Jindal, Harshita B. Saksena, Dhriti Singh, Jyothilakshmi Vadassery, Hatem Rouached, Christian Meyer, Ashverya Laxmi

## Abstract

The Target of Rapamycin Complex 1 (TORC1) is a crucial eukaryotic kinase that modulates growth in response to nutrient availability. Phosphorus (P) is an essential macronutrient, and its deficiency induces extensive reprogramming of growth and defense strategies in plants. This process involves Phosphate Starvation Response 1 (PHR1), a master regulator of the Phosphate Starvation Response (PSR). In this study, we identify a novel, non-canonical role for TORC1 in regulating P starvation responses in Arabidopsis. We demonstrate that P limitation activates TORC1, leading to the stabilization of PHR1. Inhibition of TORC1 increased sensitivity to P starvation, accompanied by disruption of starvation-induced transcriptional reprogramming. Additionally, our results reveal that the TORC1-PHR1 signaling axis plays a crucial role in reprogramming the expression of genes involved in the plant immune signaling network. This regulation is critical for the symbiotic association with the endophytic fungus *Piriformospora indica* under P starvation. These findings underscore the significant role of the TORC1-PHR1 module in orchestrating the PSR and highlight the evolutionary adaptation of TORC1 signaling pathways in plants.

## Results and Discussion

Nutrient availability, particularly the essential macronutrient phosphorus (P), is critical in shaping the growth-defense trade-offs that influence organism fitness and survival^1,2^. In *Arabidopsis thaliana*, the transcription factor Phosphate Starvation Response 1 (PHR1) functions as a master regulator of P starvation responses (PSR)^3,4^, mediating not only the plant’s adaptation to P deficiency but also its interactions with the associated microbiome during periods of nutrient limitation^1^. The Target of Rapamycin Complex 1 (TORC1) plays a pivotal role as a conserved serine/threonine protein kinase complex, acting as a central integrator of nutrient cues for growth in eukaryotes^5^. Emerging evidence suggests that TORC1 is instrumental in balancing growth and defense trade-offs in plants^6,7^, functioning as a major modulator of gene expression through interactions with various transcription factors and regulatory proteins^5,8,9^. Despite this knowledge, the potential interplay between TORC1 and PHR1 in mediating P starvation responses and plant immunity remains largely unexplored. Investigating this relationship is vital for advancing our understanding of how plants integrate nutrient sensing with growth and defense mechanisms. Such insights could have profound implications for optimizing plant resilience and productivity in nutrient-limited environments

P starvation triggers a large-scale transcriptome change to drive adaptive morphological, physiological, and defense responses^2^. To explore how P limitation affects TORC1 signaling, we first estimated the TORC1-dependent phosphorylation of S6K1 at T449 in seedlings growing in high P (HP, 625 μM P) and low P (LP, 10μM P) regimes. Remarkably, we found a significant increase in phosphorylated S6K1 levels in both the shoots and roots of seedlings growing in LP, with the effect being more pronounced in the shoots (Figure 1A). Interestingly, despite the activation of S6K1, there was no change in the phosphorylation of RPS6A (S240), a direct substrate of S6K1^10^. TORC1 phosphorylates E2Fa transcription factor to drive the expression of cell cycle genes^8^. We found no change in the expression of E2Fa and its target genes under P starvation (Figure 1B). This suggests that TORC1 signaling can take different routes, as the activation of S6K1 under P starvation didn’t trigger the canonical RPS6 and E2Fa pathways involved in promoting growth under favorable conditions. P starvation leads to enhanced promoter activity of *S6K2* in the stele and emerging lateral roots^11^. We also observed increased expression of *S6K1* and *S6K2* in P starvation (Figure 1C). However, except for a slight increase in the expression of *RAPTOR1A* and *LST8-1*, the expression levels of *TOR* and other TORC1 genes remained unchanged under P starvation.

**Figure 1.**
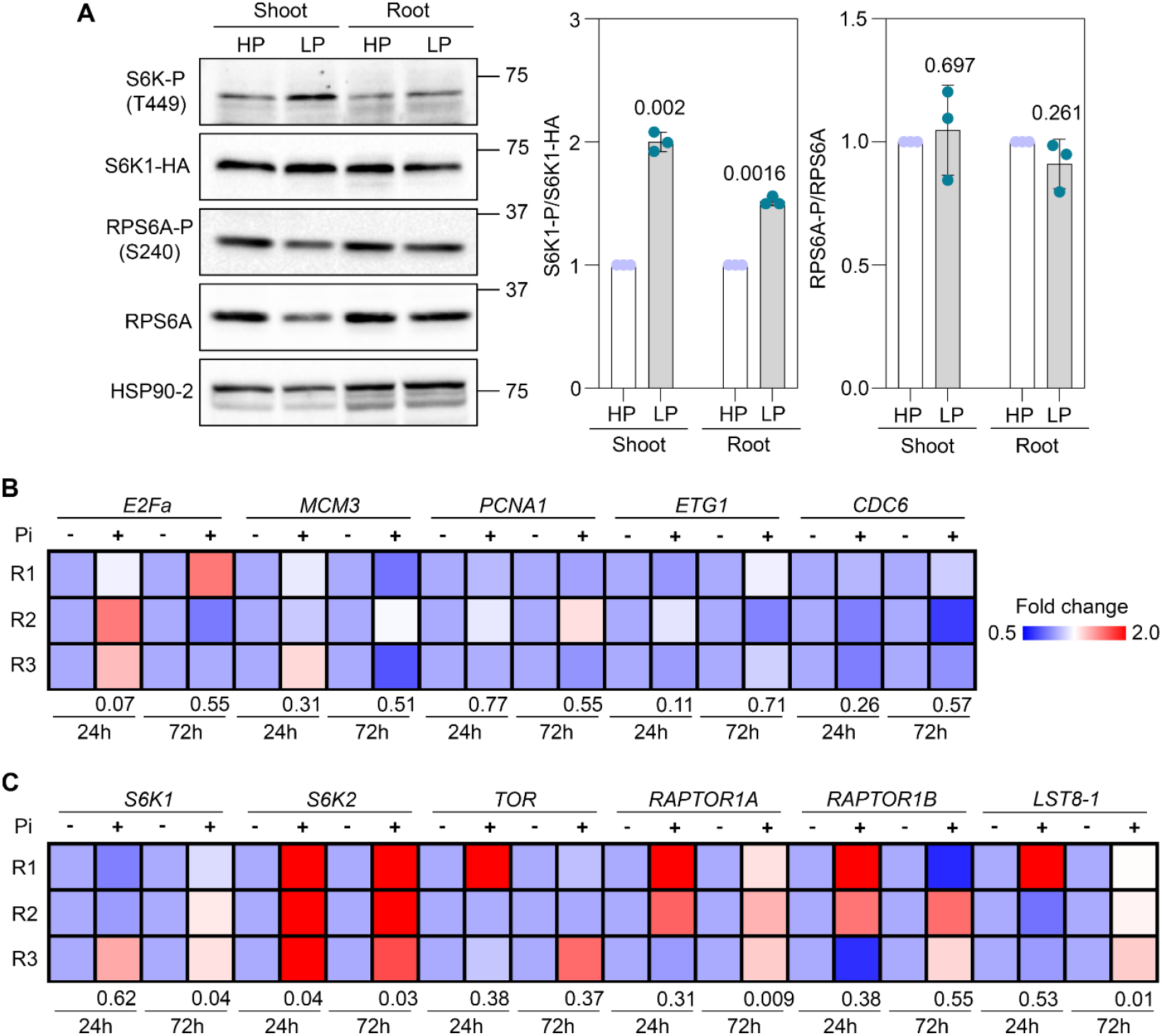
TORC1 activation during phosphate starvation. A, Estimation of TORC1 activity under high P (HP, 625 μM Pi) and low P (LP, 10 μM Pi). The TORC1-dependent phosphorylation of S6K1 (T449) and its downstream substrate RPS6A (S240) and their total levels were quantified using specific antibodies (n= 3, Student’s t-test, p-values are indicated in the graph). B, Heatmap showing relative expression of *E2Fa* and target genes in different timepoints of LP. C, Heatmap showing relative expression of *S6K1, S6K2* and TORC1 genes (*TOR, RAPTOR1A, RAPTOR1B, LST8-1*) in different time points of LP (R=replicate, n= 3, Student’s t-test, p-values are indicated below the heatmap). *UBQ10* was used as the endogenous control.

To reveal the TORC1-dependent regulation of PSR, we conducted an extensive time-dependent RNA-seq analysis (24h, 48h, and 72h) in shoot and root tissues of wild-type (WT) and *TOR* RNA interference (*tori*) mutant under high P (HP) and low P (LP) conditions (Figure 2A). We analyzed the data with WT in HP as the control to identify transcriptional changes in LP and elucidate the role of TORC1 in these conditions (Table S1). Compared to WT, we observed significant modulation of gene expression in *tori*, particularly pronounced under LP (Figure 2B). Differentially expressed genes (DEGs) formed three distinct clusters in both shoots and roots (Figure 2C, Table S2). Cluster 1 included genes typically induced in LP, such as *SPX1, IPS1*, and *SQD2*, with notably reduced induction in the *tori* mutant. This cluster encompasses mainly genes involved in PSR, and various metabolic pathways such as carboxylic acid, glucosinolate, lipid, and amino acid metabolism (Figure 2D, Table S3). Cluster 2 involved genes related to RNA metabolism and processing, translation, and primary and secondary metabolism, primarily induced in *tori* irrespective of P status. Cluster 3 comprised genes repressed in WT under long-term P starvation, including those associated with plant immunity and stress responses, many of which were induced in *tori* during HP and LP conditions. To clarify TORC1’s role in regulating PSR transcription, we analyzed the expression of 193 core PSR genes^1^, finding that their induction in LP was significantly attenuated in *tori* mutant (Figure 3A, Table S4). Given that PHR1 drives the expression of PSR genes under P deficiency, the reduced induction of several PHR1-dependent PSR genes in *tori* suggests a link between TORC1 and PHR1 (Figure S1). Further analysis of 2,364 PHR1 target genes identified via ChIP-seq^1^ revealed that a substantial subset was induced under LP conditions, particularly those in clusters 1 and 3 in shoots and clusters 1 and 2 in roots; however, activation of these clusters was largely impaired in *tori* (Figure 3B, Table S5). These clusters mainly include genes involved in PSR, P and carboxylic acid metabolism (Figure S2, Table S6). Notably, genes involved in plant immunity and stress responses repressed in wild-type plants during long-term starvation (cluster 5 in shoots and cluster 4 in roots), displayed no such repression in *tori* under LP (Figure 3B, Figure S2, Table S6). Supporting these findings, we observed impaired induction of several PSR marker genes in β-Estradiol-induced TOR RNAi (*tor-es1*) seedlings following TOR inhibition (Figure 3C), collectively indicating a crucial role of TORC1 in modulating PHR1 function and the overall regulation of plant responses to P starvation.

**Figure 2.**
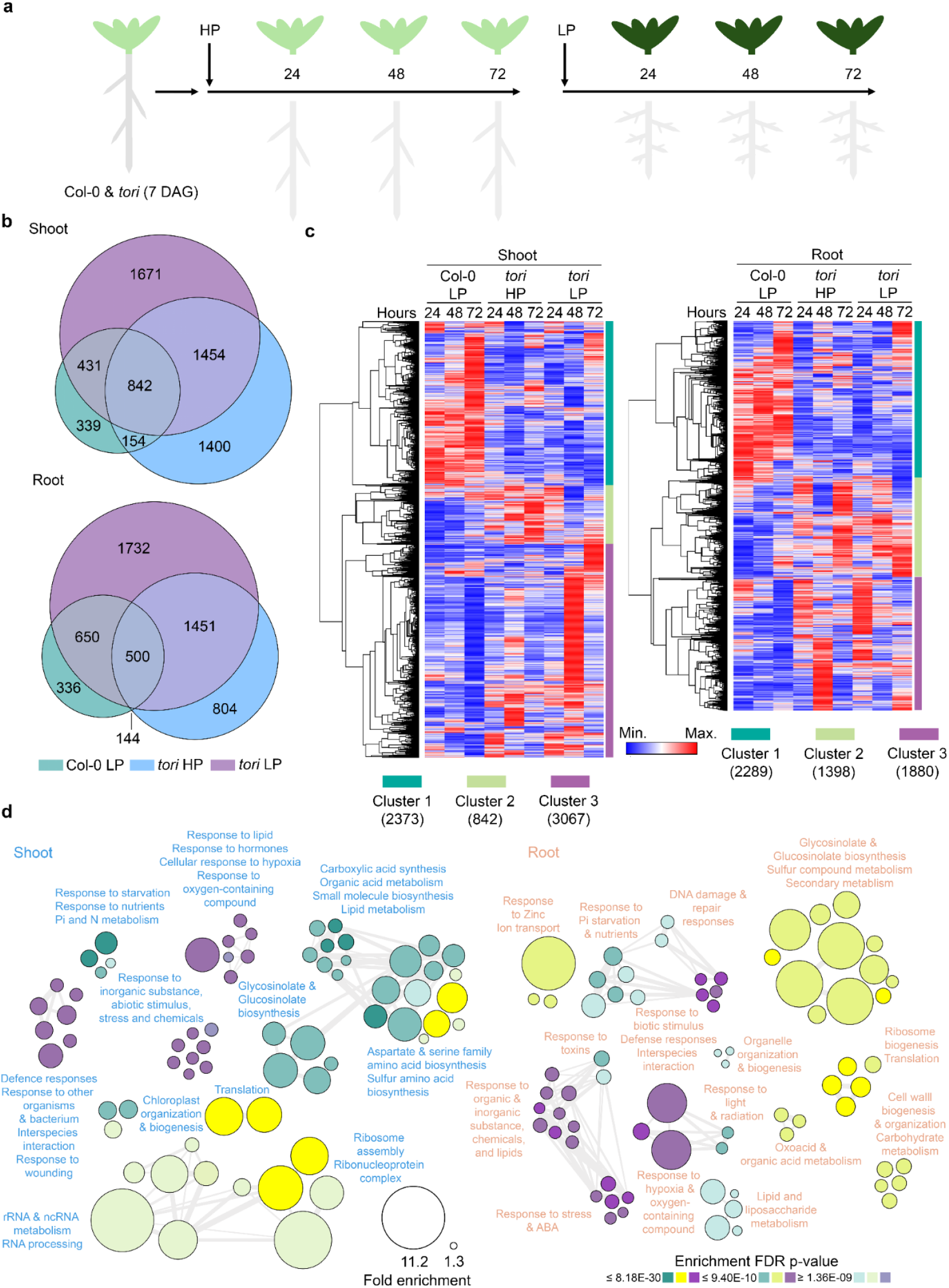
TORC1 is important for the transcriptional reprogramming during phosphate starvation. A, Graphical representation of experimental setup used for RNA-seq sample collection. B, Euler diagram showing the overlap in differentially expressed genes (DEGs) in shoot and root. C, Heatmap showing different clusters of DEGs. Clustering was performed by One minus Pearson’s correlation with average linkage method. Number of genes in each cluster is shown in brackets. D, Gene ontology (biological process) network of DEGs. The size of the bubble represents fold enrichment. The color represent cluster and color intensity represent fold enrichment FDR p-value.

**Figure 3.**
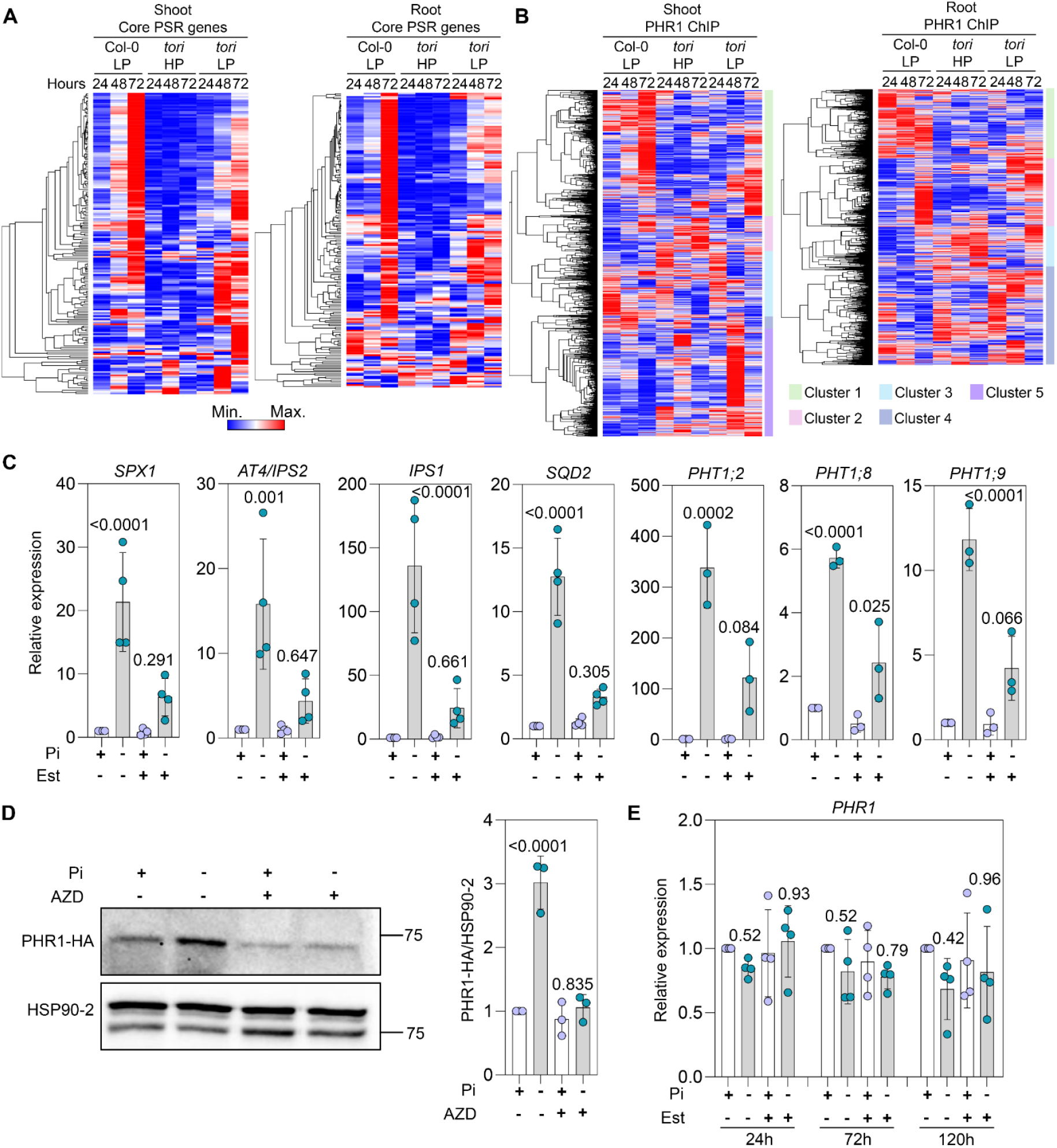
TORC1 drives the expression of phosphate starvation response genes by promoting PHR1 stability. A, Heatmap showing the expression pattern of curated 193 core PSR genes in Col-0 and *tori* shoot and root tissues. B, Heatmap showing the expression pattern of 2364 PHR1 targets (Identified by ChIP-seq) in Col-0 and *tori* shoot and root tissues. Clustering was performed by One minus Pearson’s correlation with average linkage method. C, Expression of PSR marker genes *tor-es1* in high (+, 625 μM Pi) and low (-, 10 μM Pi) P with and without TOR inhibition using 10 μM β –Estradiol (n=3/4, One-way ANOVA with Tukey’s multiple comparisons test, p-values are indicated in the graphs). D, The level of PHR1-HA in *UBQ10:PHR1-HA* line grown in high and low P regimes for 72h with and without TOR inhibition using 1 μM AZD8055. The level of PHR1-HA is quantified relative to loading control HSP90-2 (n=3, One-way ANOVA with Tukey’s multiple comparisons test, p-values are indicated in the graphs). E, The expression of *PHR1* in high and low P regimes with and without TOR inhibition using 10 μM β –Estradiol (n=3, One-way ANOVA with Tukey’s multiple comparisons test, p-values are indicated in the graphs). *UBQ10* was used as the endogenous control for RT-qPCR experiments.

PHR1 stability is regulated by ubiquitination, with P starvation promoting its stability^12^. To investigate the role of TORC1 in regulating PHR1 stability, a *UBQ10:PHR1-HA* line was grown in HP and LP conditions with or without TOR inhibition using AZD8055 (Figure 3D). As reported^12^, the PHR1 level was increased in LP. However, this increase was abolished by TOR inhibition. Thus, TORC1 is crucial for enhancing PHR1 stability during P starvation to drive the expression of PSR genes. Nonetheless, no effect of TORC1 was found on the expression of PHR1 in HP and LP conditions (Figure 3E). Thus, TORC1 regulates PHR1 function by enhancing its stability during P starvation, thereby fine-tuning the downstream events controlled by PHR1. In addition to activating PSR gene expression, PHR1 also directly binds to the promoters of plant immunity genes to suppress their expression. This regulation is crucial for shaping the microbiome and interacting with beneficial microbes in P starvation^1^. Loss of *tori* leads to the induction of immunity signaling genes especially in LP (Figure 2C, 2D). To explore this further, we analyzed the expression of plant defense hormone-(Jasmonic acid and Salicylic acid) and flg22-induced genes in our dataset (Figure 4A, 4B, Table S7, S8). Notably, a significant number of these immunity marker genes are induced in LP (Cluster 1 & 2 in shoot and root). This includes genes related to PSR, plant defense, and various metabolic pathways, including primary (Pi, lipid, and organic acid) and secondary (glucosinolate and glycosinolate) metabolism (Figure S3, S4, Table S9, S10). The induction of these genes was impaired in *tori*. A large subset of immunity marker genes was largely repressed in LP, especially in the shoot at 72h (Clusters 3 & 4 in JA/SA, Cluster 3 in flg22). This includes core plant immunity genes such as *CBP60g, WRKY70, JAZs (JAZ1, JAZ3, JAZ5, JAZ6, JAZ7, JAZ10, JAZ12)* and *LOX3*. Repression of these clusters was notably reduced in *tori* (Figure 4A, 4B, Table S7-S10). Interestingly, the *phr1 phl1* mutant showed increased expression of plant defense genes, independent of P status^1^. Similarly, many core defense signaling genes in *tor-es1* mutants followed this pattern (Figure 4C).

**Figure 4.**
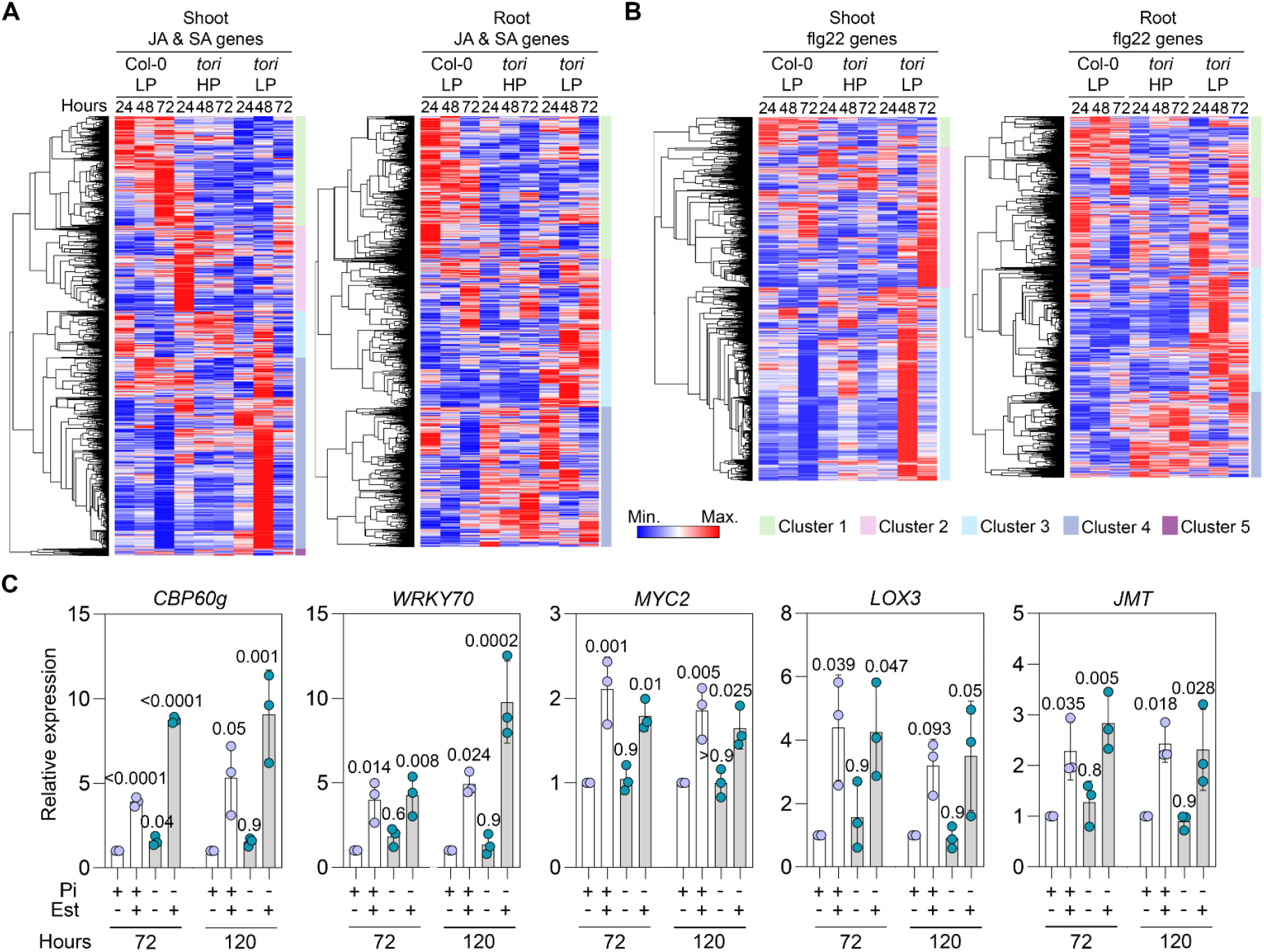
TORC1 modulates immune signaling during phosphate starvation. A, Heatmap showing the expression pattern of 3029 JA/SA marker genes (induced by JA/SA) in Col-0 and *tori* shoot and root tissues. B, Heatmap showing the expression pattern of 2690 flg22 marker genes (induced by flg22) in Col-0 and *tori* shoot and root tissues. Clustering was performed by One minus Pearson’s correlation with average linkage method. C, Expression of plant immunity marker genes *tor-es1* in high (+, 625 μM Pi) and low (-, 10 μM Pi) P with and without TOR inhibition using 10 μM β –Estradiol (n=3, One-way ANOVA with Tukey’s multiple comparisons test, p-values are indicated in the graphs). *UBQ10* was used as the endogenous control for RT-qPCR experiments.

Collectively, these results indicate that TORC1 is a crucial regulator of gene expression in P starvation through the PHR1 signaling. Loss of TORC1 leads to failure in inducing PSR gene expression and modulating plant immunity signaling network. To validate the role of TORC1 in the adaptive strategies during P starvation, we analyzed the wild-type (WT) plants and *TOR* RNA interference line (*tori*) under HP and two LP (100 μM/moderate starvation and 10 μM/severe starvation) conditions. Under HP conditions, there was no significant difference in primary root (PR) length between WT and *tori* plants (Figure 5A). However, P starvation reduced PR length in both genotypes, with a much more severe reduction observed in *tori* under moderate and severe P limitation. Consistent with this, the root apical meristem (RAM) in *tori* was smaller and showed a higher degree of terminal differentiation under P starvation (Figure 5B). We then tested a TOR overexpression line (*TOR OE1*) under these conditions. The *TOR OE1* plants were less sensitive to P starvation, exhibiting significantly longer PR length and larger RAM size than WT under P starvation (Figure S5). These results confirm the role of TORC1 as a critical factor in determining the sensitivity of seedlings to P starvation. Changes in internal P levels can influence the sensitivity to P starvation. Therefore, we measured the levels of both soluble and total P in WT and *tori*. P starvation reduced the level of soluble and total P; however, no difference in the P level was observed between WT and *tori* in both HP and LP conditions (Figure S6). Furthermore, while the deposition of iron (Fe) in the RAM apoplast is known to be associated with the inhibition of RAM activity during P starvation^13^, the PR growth inhibition in P starvation was rescued similarly in both WT and *tori* in the absence of Fe (Figure S7). This result indicates that the increased sensitivity of *tori* to P starvation is not due to changes in overall P level or Fe-dependent signaling in the root.

**Figure 5.**
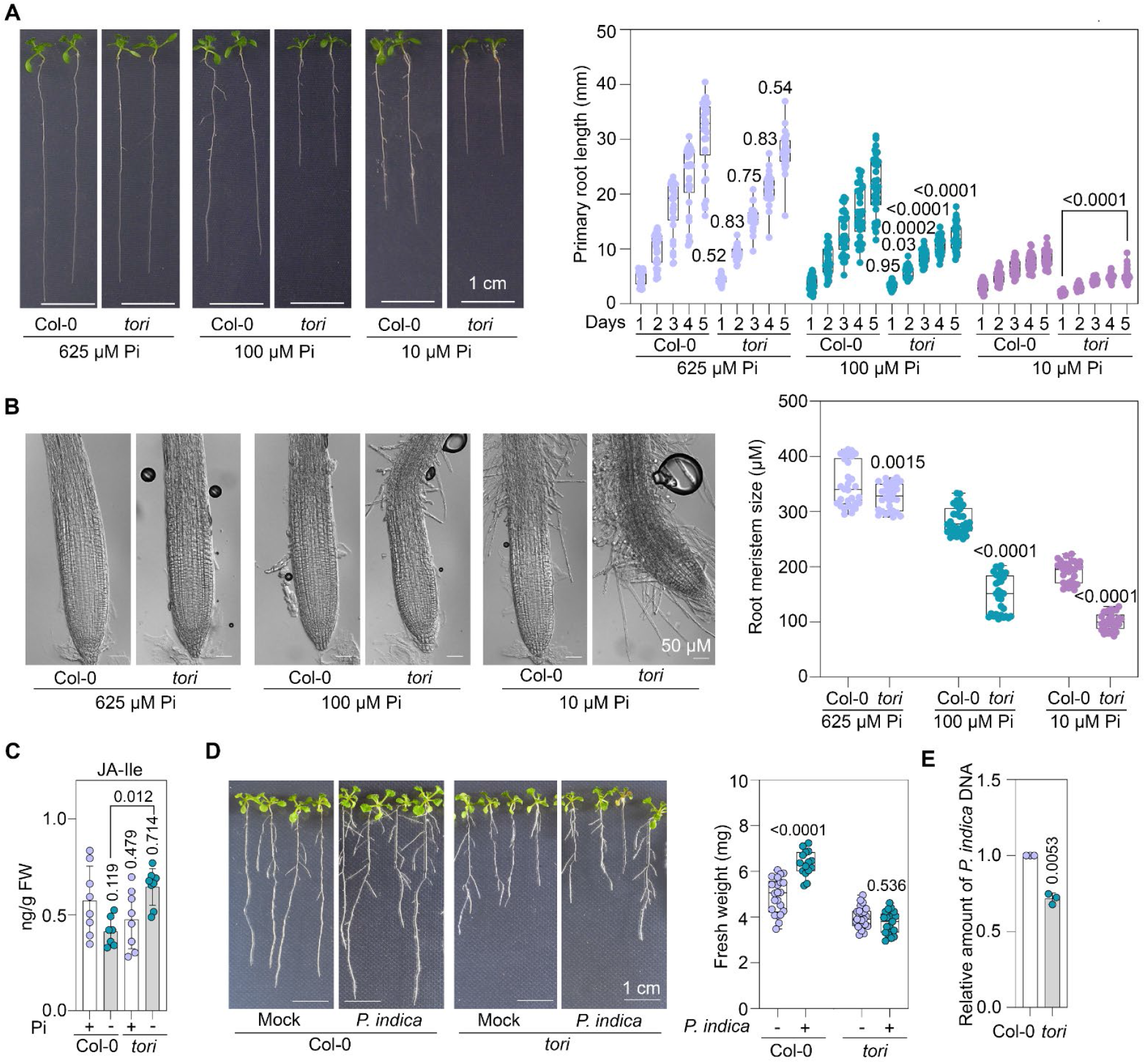
TORC1 is critical for the tolerance in phosphate starvation and association with the beneficial endophyte. A, Phenotype and primary root growth kinetics of Col-0 and *tori* in HP and under different LP conditions. B, Root meristem size of Col-0 and *tori* in HP and under different LP regimes. Experiments were repeated three times, and the graphs indicate a biological replicate (n=30, two-way ANOVA with Tukey’s multiple comparisons test, p-values are indicated in the graphs) was performed between Col-0 and *tori* growing in same Pi regimes, p-values are indicated in the graphs). C, Estimation of JA-Ile level in Col-0 and *tori* in HP and LP regimes (n=7-10, One-way ANOVA with Tukey’s multiple comparisons test, p-values are indicated in the graphs). D, Phenotype and biomass of Col-0 and *tori* growing in low P with or without the beneficial fungi *Piriformospora indica* (n=15-23, One-way ANOVA with Tukey’s multiple comparisons test, p-values are indicated in the graphs). E, Level of *P. indica* DNA in Col-0 and *tori* growing in low P (n= 3, Student’s t-test, p-values are indicated in the graph).

The intricate balance between nutrient availability and plant immunity emerges as a crucial factor in plant responses to P starvation^2^. Transcriptome analysis identified a major role of TORC1 in regulating JA, SA, and ABA signaling networks during P starvation. Quantification of these hormones identified a higher level of bioactive JA (JA-Ile) in *tori* specifically in LP (Figure 5C, Figure S8). The PHR1-dependent modulation of plant immunity is a critical factor regulating the microbial association required for mitigating P starvation^1^. *Piriformospora indica* is an endophytic fungus that helps plants alleviate P starvation^14^. The JA and immunity signaling pathways are crucial for maintaining a mutualistic association with *P. indica*^15^. In WT, co-cultivation of *P. indica* reduced the negative effect of P starvation on plant growth (Figure 5D). However, this effect was not observed in *tori* seedlings. Supporting this, *tori* displayed reduced colonization by *P. indica* (Figure 5E). This is in line with the higher level of JA-Ile and misregulation of plant immunity genes in *tori* (Figure 4A, 4B, 5C). Thus, the dysregulation of the plant immunity signaling networks in *tori* affects the microbial association important for alleviating the negative effects of P starvation.

Our findings reveal a crucial role for TORC1 in regulating PHR1 signaling during P starvation in Arabidopsis. This regulation is pivotal for driving the expression of PSR genes and modulating the immunity signaling network and the association with the beneficial microbe, *P. Indica*. Interestingly, the activation of TORC1 signaling under P starvation diverges from its canonical role in promoting growth-related pathways like the RPS6 and E2Fa signaling axes, which are important for promoting translation, cell division, and general growth under favorable conditions^5^. Instead, during P starvation, TORC1 activation enhances PHR1 stability and signaling. PHR1 is regulated by post-translational modifications such as phosphorylation, ubiquitination, and SUMOylation^12,16–18^. SnRK1, the antagonistic kinase of TORC1, phosphorylates PHR1, inhibiting its ability to activate gene expression^16^. However, the mechanism by which TORC1 affects the stability of PHR1 is still unclear. This regulation may involve an interaction between TORC1 and SnRK1, direct phosphorylation by TORC1, or the involvement of downstream signaling components like S6Ks. Our findings represent a significant change in the understanding of plant TORC1 signaling, as studies primarily identified its role as a driver of growth in favorable conditions. TORC1 is important for promoting growth under P-abundant conditions in Arabidopsis^19^. It is possible that depending on the signal, different TORC1 signaling circuits in plants execute specific functions in the cell. For instance, sulfur limitation triggers contrasting signaling responses in root and shoot tissues by modulating the TORC1 signaling, indicating the existence of distinct TORC1 signaling circuits in plants as well^20^. In yeast and mammals, spatially distinct pools of TORC1 at the cellular level modulate different cellular processes^21,22^. Our findings demonstrate an interplay between TORC1 and PHR1, revealing the integration of TORC1-PHR1 axis nutrient sensing and defense in plants.

## Acknowledgments

We acknowledge Confocal and Metabolomics facilities for their assistance and DBT-eLibrary Consortium (DeLCON) for providing access to e-resources.

## Supplemental information

Figure S1-S8

Table S1-S11

## Author contributions

Conceptualization, A.L., and M.J.K.; methodology, A.L., M.J.K., P.A., J.V., and M.K.; formal analysis, M.J.K., P.A., M.K., and J.V.; investigation, P.A., M.J.K., M.K., S.M., A.T., S.J., H.B.S., D.S., M.S., and M.S.; data curation, P.A., M.J.K., J.V., and A.L.; writing – original draft preparation, M.J.K., P.A., and H.R.; writing – review & editing, A.L., J.V., and C.M.; visualization, M.J.K.; supervision, A.L., and M.J.K.; project administration, A.L., and M.J.K., Funding acquisition, A.L., and M.J.K.

## Data and code availability

Transcriptome raw data will be deposited to NCBI’s Sequence Read Archive (SRA). All data generated in this study are available in the figures, tables and data files associated with this manuscript. Additional information and requests for resources generated in this study are available from the lead contact, Ashverya Laxmi.

## Materials and Methods

### Plant material and growth conditions

Arabidopsis Columbia (Col-0) ecotype was used as wildtype (WT) in this study. The seeds of *tor-es1*^*23*^ (CS69829) and 35S::*S6K1-HA*^*23*^ (CS73259) were procured from ABRC. The UBQ::*PHR1-HA*^*12*^ seeds were provided by Su-Hyun Park. The *TOR OE1* (*GK-166C06*) and *TOR 35-7* (*tori)* lines are described in a previous study^*24*^. Seeds were surface sterilized using 20% (v/v) commercial bleach and 0.001% SDS for 10 minutes and subsequently washed five times with sterilized Milli-Q water. Sterilized seeds were kept at 4°C for 48 h. Seeds were plated on 0.5X Murashige & Skoog (MS) medium (pH: 5.7) with 0.8% agar and 30 mM sucrose (Sigma-Aldrich) in square Petri plates. For the experiments in different P availability, 9 mM sucrose was used as sugars affect PSR gene expression^25^. Seedlings were grown vertically in a tissue culture room under controlled conditions (16 h light: 8 h dark photoperiod at 22°C ±2 and average photosynthetically active radiation of 60 μmol m-2 s-1). Adult plants were grown in the same conditions in pots containing agro peat and vermiculite (3:1).

### Phenotypic analysis

Sterilized seeds were plated on 0.5X MS solid medium and grown under controlled conditions in square Petri plates. For the phenotypic analysis in P starvation, 7 days-old uniformly grown seedlings were transferred to 0.5X MS medium with modified inorganic phosphate (Pi) concentration of 625 μM of KH_2_PO_4_ (Sigma-Aldrich) as high, 100 μM as moderate, and 10 μM as severe low P medium. Primary root length was marked at the beginning of the experiment and on every day for 5 days. For Fe-P phenotypic analysis, 7 days-old uniformly grown seedlings were transferred to P rich and deficient medium in combination with the 50 μM and 0 μM of FeSO_4_ (Sigma-Aldrich). Primary root length was marked after the transfer. The Petri plates were photographed using a Nikon camera and primary root length was quantified using ImageJ (NIH, USA). For the RAM length measurement, seedlings growing in different P regimes for 5 days were treated with clearing solution (8:3:1 ratio of chloral hydrate: water: glycerol) for 2 min. The images of roots were taken by TCS SP5 microscope (Leica Microsystems).

### Soluble and total phosphate estimation

The ascorbate-molybdate assay was used for soluble P estimation^26^. Seedlings grown in 0.5X MS solid medium in standard controlled conditions for 7 days were transferred to 0.5X MS solid medium with 625 μM and 10 μM of Pi for 5 days. Approximately 40 mg of seedlings were lysed using Tissuelyser II (Qiagen) in liquid nitrogen. Samples were resuspended in 1% glacial acetic acid and centrifuged at 13000 rpm for 10 minutes. The supernatant was dissolved in an equal volume of phosphomolybdate reagent (1:6 ratio of ascorbic acid and ammonium molybdate). Absorbance was measured at 820 nm using a POLARstar Omega plate reader. The P level was quantified using a standard curve and normalized per gram of fresh weight. Similar treatment was done for total P estimation. 150 mg of seedlings were oven-dried and proceeded to the muffle furnace for ashing at 550°C for 5 h. Samples were re-suspended in 2 mL of 2 N HCl for 20 min and centrifuged at 13000 rpm for 5 minutes. The supernatant was mixed with an equal volume of vanadate-molybdate reagent (Sigma-Aldrich). The absorbance of the samples in 96 welled plate was taken using POLARstar Omega plate reader at 410 nm.

### Sample preparation and RNA-seq analysis

Col-0 and *tori* seedlings were grown in 0.5X MS solid medium in standard controlled conditions for 7 days. These seedlings were transferred to 0.5X MS solid medium with 625 μM and 10 μM of Pi in Petri plates. The root and shoot samples were collected after 24, 48, and 72 h of treatments. The samples were harvested in liquid nitrogen and stored at -80°C. Total RNA was extracted using the RNeasy® plant mini kit (Qiagen). The RNA was purified using the sodium acetate-ethanol method. The quantity and quality of RNA was identified using Agilent 2100 Bioanalyzer (Agilent Technologies) respectively. RNA-seq library was prepared using NEB Next Ultra II RNA library prep kit for Illumina (New England Biolabs) according to the manufacturer’s protocol. Quantity and quality of cDNA libraries were assessed using Qubit dsDNA HS Assay Kit (Thermo Fisher Scientific), Agilent 2100 Bioanalyzer and by visualization on gel using SYBR™ Gold nucleic acid gel stain (Thermo Fisher Scientific). The 150 bp paired-end sequencing was performed on the Illumina HiSeq X Ten platform. QC was performed using FastQC following the removal of the adapter sequence using trimmomatic. The sequencing reads were mapped to the reference Arabidopsis thaliana genome (TAIR10) using HISAT2 in default parameters. The DEGs were identified using DESeq2 in the default parameter. The DEGs in at least one condition/genotype (FDR p-value of <0.05) were used for further analysis. The list of the core PSR genes, PHR1 ChIP-seq targets, JA/SA and flg22-responsive genes were retrieved from a previous study^1^. The relative heatmaps and clustering was performed in Morpheus (https://software.broadinstitute.org/morpheus/). The clustering of DEGs were performed by One minus Pearson’s correlation with average linkage method. The GO analysis was performed using ShinyGo v0.80. The GO data was imported in REVIGO v1.8.1 and visualized in Cytoscape 3.10.2.

### Sample preparation and western blotting

The 35S::*S6K1-HA* seedlings were used for the detection of S6K1-T449, S6K1-HA, RPS6A-S240, RPS6A estimation under different P conditions. The 7 DAG seedlings growing in standard growth conditions were transferred to 0.5X MS solid medium with 625 μM and 10 μM of Pi in a Petri plate under controlled conditions for 96h. The root and shoot tissues were excised and harvested in liquid nitrogen. Samples were stored at -80°C. Similarly, the UBQ10:*PHR1-HA* seedlings growing in standard growth conditions were transferred to 0.5X MS liquid medium with 625 μM and 10 μM of Pi with or without 1 μM of AZD8055 (MedChemExpress) for three days. Samples were harvested in liquid nitrogen and stored at -80°C.

Frozen samples were lysed using Tissuelyser II (Qiagen) in liquid nitrogen. Crushed samples were re-suspended in ice-cold protein extraction buffer (137 mm NaCl, 4.3 mm Na_2_HPO_4_, 2.7 mm KCl, 1.47 mm KH_2_PO_4_, 10% glycerol, 1/500 (v/v) plant-specific protease inhibitor cocktail, 1/500 (v/v) phosphatase inhibitor cocktail 3) in ice. The soluble protein was extracted by centrifugation at 13000 rpm at 4°C. Protein concentration was quantified using Quick start Bradford reagent (Bio-rad) using 96 well plate in POLARstar Omega plate reader (BMG Labtech). An equal amount of protein was boiled with SDS Laemmli buffer at 95°C and was separated on 12.5% SDS gel. Separated proteins were transferred to the nitrocellulose membrane (GE healthcare). The membrane was incubated with 0.5% w/v BSA prepared in 1X TBST (Tris-buffered saline with 0.01% v/v Tween 20) buffer for 1 h. Membrane was further incubated in the antibodies diluted with 1X TBST solution for overnight. For the detection of total PHR1-HA and total S6K-HA, an antibody against HA tag (Anti-HA tag, dilution: 1:2500, catalog no.: 2367, CST) was used. To detect the phosphorylated form of S6K, S6Kp antibody (Phospho-p70 S6 kinase pThr389 antibody; dilution: 1:2000; catalog no.: MA5-15117, Invitrogen) was used. HSP90-2 antibody (Heat shock protein 90-2 antibody; dilution: 1:10000; catalog no.: AS11 1629, Agrisera) was used as the loading control. Biorad (Clarity ECL substrate) developer Solution was used to develop luminescence and visualized by Chemidoc XRS+ imaging system (Bio-rad). After the immunodetection, membranes were stripped using mild glycine buffer (200 mM glycine, 3.5 mM SDS, 0.1% v/v Tween 20, pH:2.2) after washing with 1X TBST buffer, stripped membranes were again blocked with 0.5% w/v BSA and proceeded for immunodetection. Blots were quantified using ImageJ (NIH).

### Treatments and qRT-PCR analysis

For the gene expression analysis in *tor-es1*, seedlings were grown on 0.5X MS solid medium for 7 days. The seedlings were transferred to 0.5X MS liquid medium with 625 μM and 10 μM of Pi with and without 10 μM of β-Estradiol (Sigma) in a 6-welled plate and incubated at 22°C on an orbital shaker (100 rpm) for 24h. For long-term P deficiency treatments, *tores1* seedlings were transferred to 0.5X MS solid medium with 625 μM and 10 μM of Pi with and without 10 μM of β-Estradiol in Petri plates. All samples were harvested in liquid nitrogen and stored at -80°C. Same protocol was used for gene expression analysis in Col-0.

The frozen samples were lysed in liquid nitrogen using Tissuelyser II (Qiagen) with tungsten beads. Total RNA was isolated using the RNeasy® plant mini kit (Qiagen) as instructed in the user manual. NanoDrop 2000 spectrophotometer (Thermo Fisher Scientific) was used for the quantification of RNA. An equal quantity of RNA was used to prepare cDNA using a High-Capacity cDNA reverse transcription kit (Thermo Fisher Scientific). Diluted cDNA samples (1:20) were used for quantitative real-time PCR (qRT-PCR) using SYBR green (TB GREEN PREMIX). The reaction was carried out in 96/384-well optical reaction plates (Applied Biosystems) using Applied Biosystems real-time PCR systems. Primers for qRT-PCR were designed on Primer Express v3.0 (Thermo Fisher Scientific). The delta-delta CT method was used to calculate the mRNA level ^27^. The *UBIQUITIN 10 (UBQ10)* gene was used as the endogenous control. The primers used are listed in Table S11.

### Hormone estimation

Phytohormones were quantified as described earlier^28^. Seedlings grown on 0.5X MS solid medium were transferred to 0.5X MS solid medium with 625 μM and 100 μM of Pi in a Petri plate and grown under controlled conditions for 8 days. 250 mg of seedling fresh weight was harvested using liquid nitrogen and stored at -80°C. Frozen samples were ground into a fine powder using a Tissuelyser II. Samples were dissolved using 1.5 mL of ice-cold methanol containing 40 ng of d6-JA (HPC Standards), 40 ng of salicylic acid-d4 (HPC Standards), 40 ng of d6-ABA (HPC Standards) and 8 ng of d6-JA-Ile conjugate (HPC Standards) as labeled internal standards. Dissolved samples were mixed using a rotator shaker followed by centrifugation at 13000 rpm for 30 mins. Samples were vacuum-dried for 2 h at 10000 rpm and quickly dissolved in 0.5 mL of ice-cold methanol. A triple-quadruple LC-MS/MS system (QTRAP 6500+) was used for phytohormone quantification.

### *Piriformospora indica* co-cultivation and DNA quantification

The seedlings were grown on 0.5X MS solid medium under controlled conditions in square Petri plates for 7 days. *P. indica* was cultured on Kaefer medium at 28°C in dark for 21 days. Small discs of *P. indica* were cut using cork-borer and used for co-cultivation experiments. At 7^th^ day, seedlings were transferred to 0.5X MS medium with 625 μM and 100 μM of Pi containing control and *P. indica* discs. The Petri plates were kept for 8 days in the standard growth conditions. Seedlings were photographed using a Nikon camera and fresh weight of seedlings was measured using a microbalance (Sartorius, Germany). For the *P. indica* DNA quantification, roots were harvested on the 8th day of co-cultivation in liquid nitrogen. DNA was extracted using the CTAB extraction buffer^29^. An equal amount of root DNA was used for qRT-PCR analysis. The *ITS* gene of *P. indica* was used for the quantification of *P. indica* colonization while *UBQ10* of Arabidopsis was used as normalization control. Relative change in fungal DNA was calculated using the delta-delta CT method. The primers used are listed in Table S11.

### Statistical analysis

The number of biological replicates (n) and the statistical analysis used for the experiment is mentioned in figure legends. All experiments were performed at least three times. GraphPad Prism v.10 and Microsoft Excel were used for graph preparation and statistical analysis.

---

**Figure S1.**
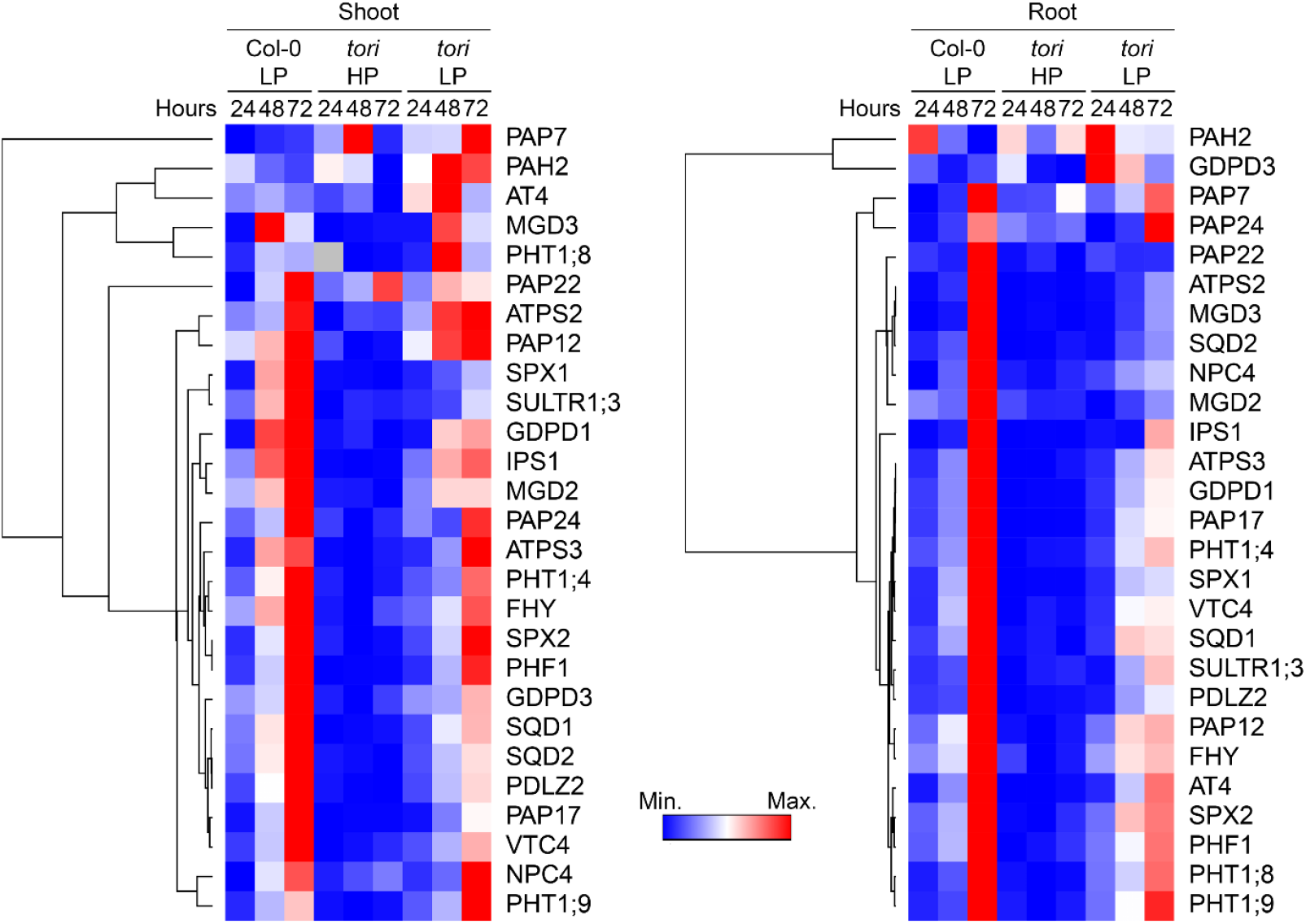
TOR is important for inducing PHR1-dependent phosphate starvation response genes. Heatmap showing the expression pattern of 27 PHR1-dependent PSR genes in Col-0 and *tori* shoot and root tissues. Clustering was performed by One minus Pearson’s correlation with average linkage method.

**Figure S2.**
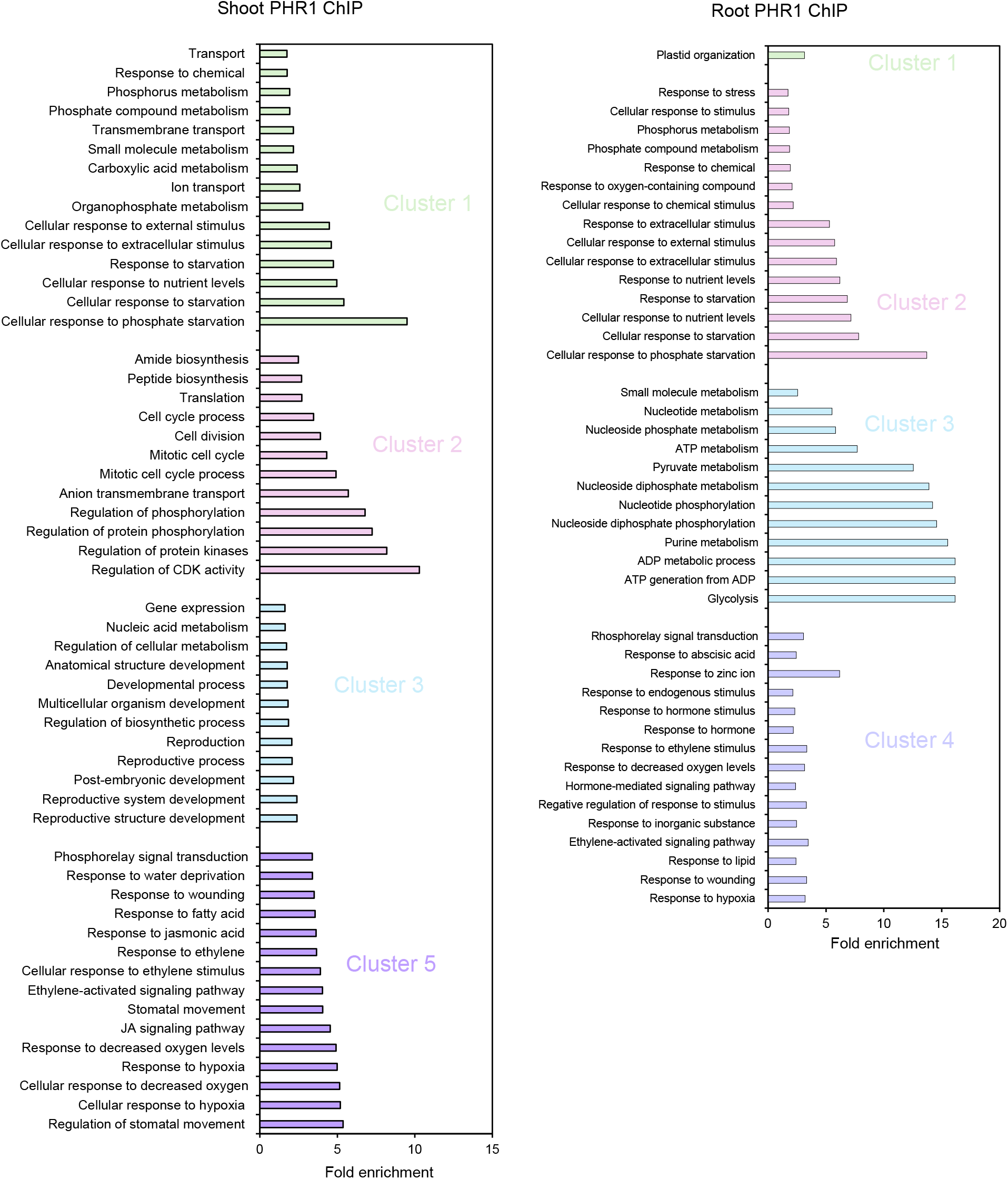
Biological processes enriched in different clusters of PHR1 target genes. Gene ontology chart of biological processes overrepresented in different clusters of PHR1 target genes (Identified by ChIP-seq). The clusters are represented in Figure 3B.

**Figure S3.**
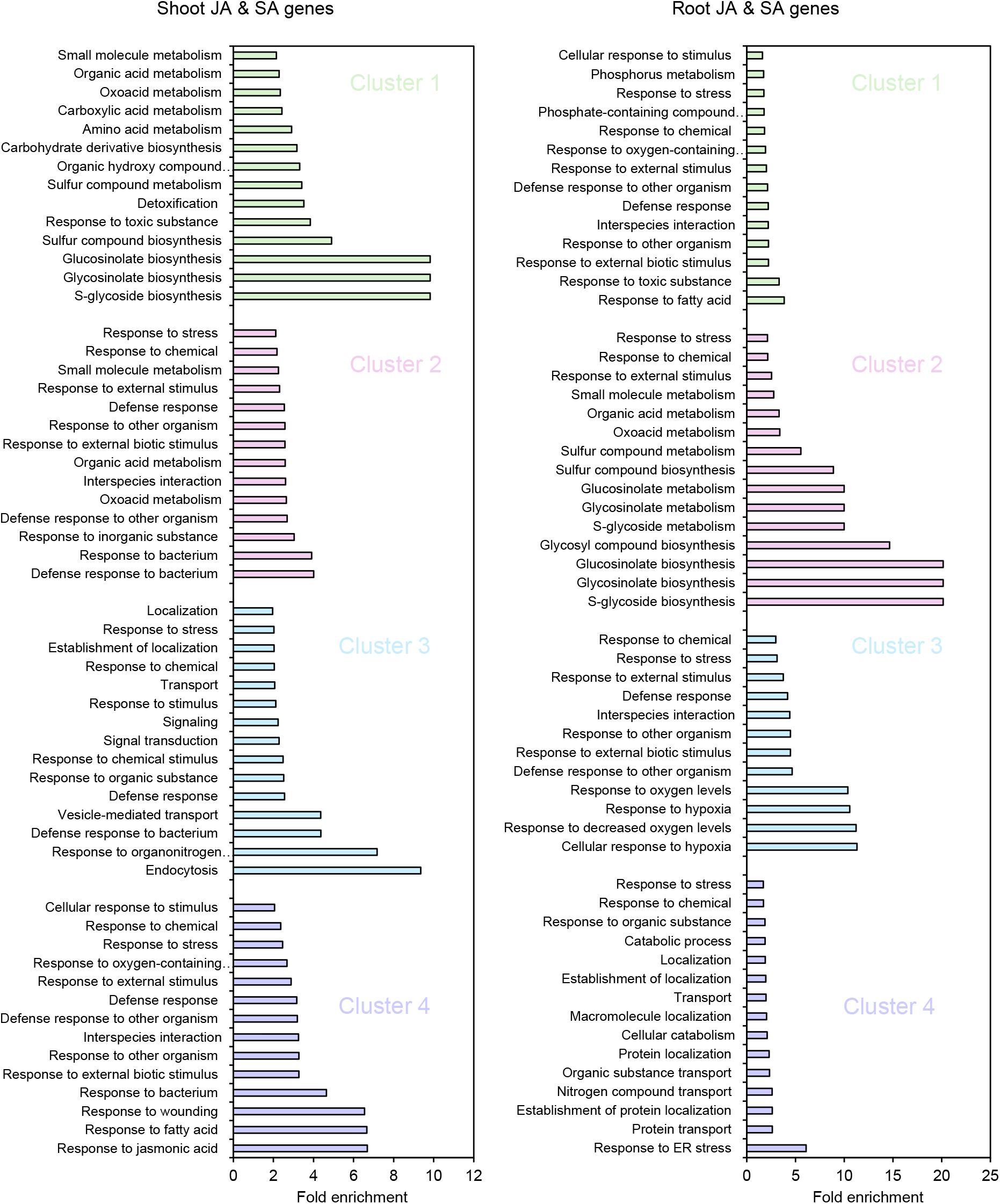
Biological processes enriched in different clusters of JA & SA marker genes. Gene ontology chart of biological processes overrepresented in different clusters of JA & SA marker genes. The clusters are represented in Figure 4A.

**Figure S4.**
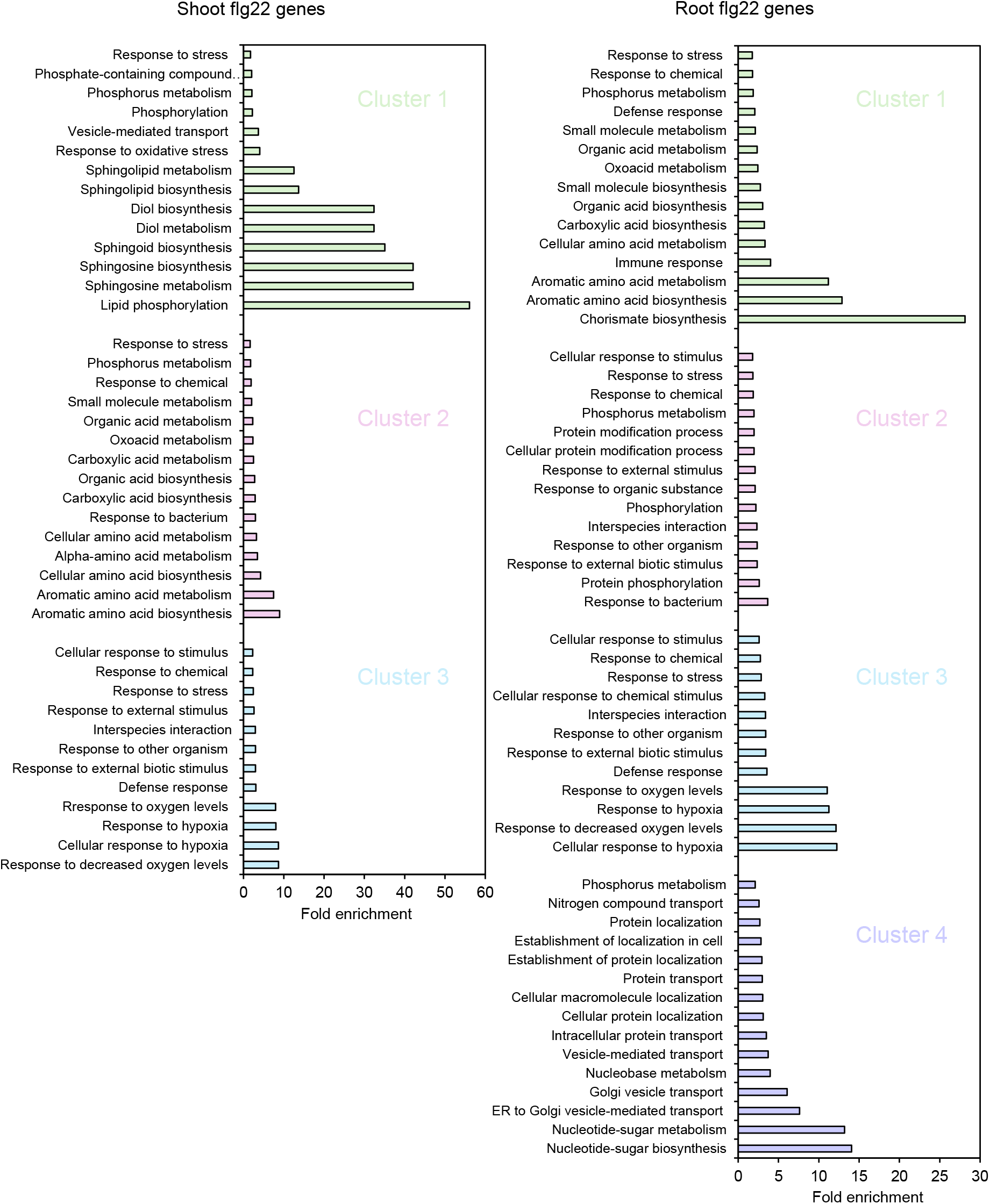
Biological processes enriched in different clusters of flg22 marker genes. Gene ontology chart of biological processes overrepresented in different clusters of flg22 marker genes. The clusters are represented in Figure 4B.

**Figure S5.**
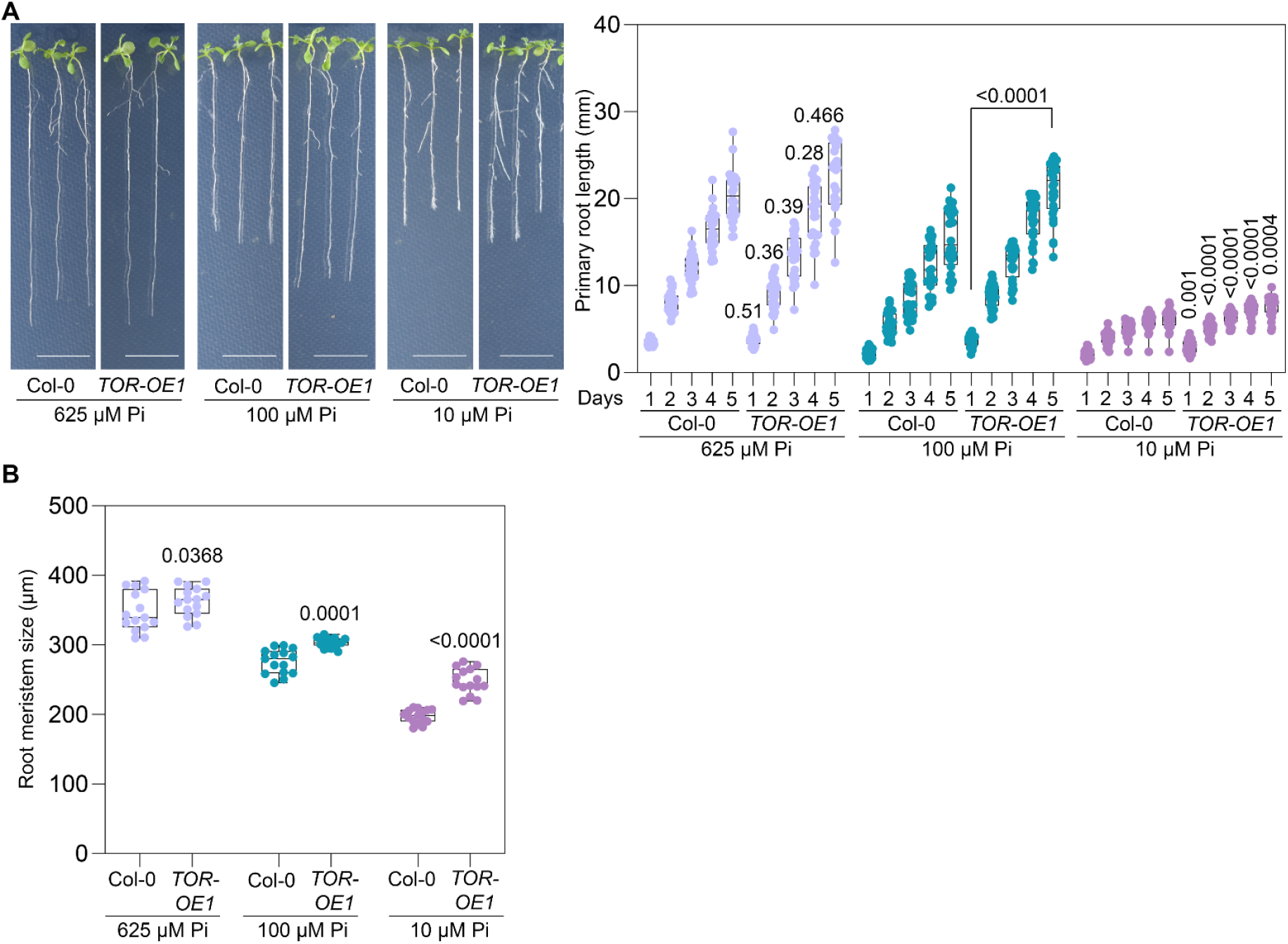
TOR overexpression leads to enhanced resistance to phosphate starvation. A, Phenotype and primary root growth kinetics of Col-0 and *TOR-OE1* in HP and under different P limitation conditions (n=28, Statistical analysis (two-way ANOVA with Tukey’s multiple comparisons test) was performed between Col-0 and *TOR-OE1* growing in same P regimes, p-values are indicated in the graphs). B, Root meristem size of Col-0 and *TOR-OE1* in HP and under different P regimes. Experiments were repeated three times, and the graphs indicate a biological replicate (n=15, Statistical analysis (two-way ANOVA with Tukey’s multiple comparisons test) was performed between Col-0 and *TOR-OE1* growing in same P regimes, p-values are indicated in the graphs).

**Figure S6.**
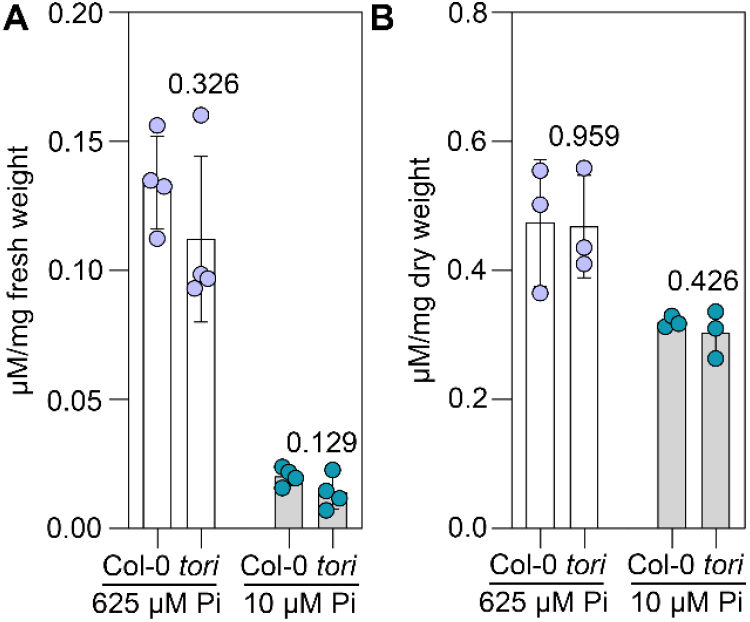
Soluble and total P in different regimes. A, Level of soluble P in Col-0 and *tori* growing in different P regimes (n= 4, Student’s t-test, p-values are indicated in the graph). B, Level of total P in Col-0 and *tori* growing in different P regimes (n= 3, Student’s t-test, p-values are indicated in the graph).

**Figure S7.**
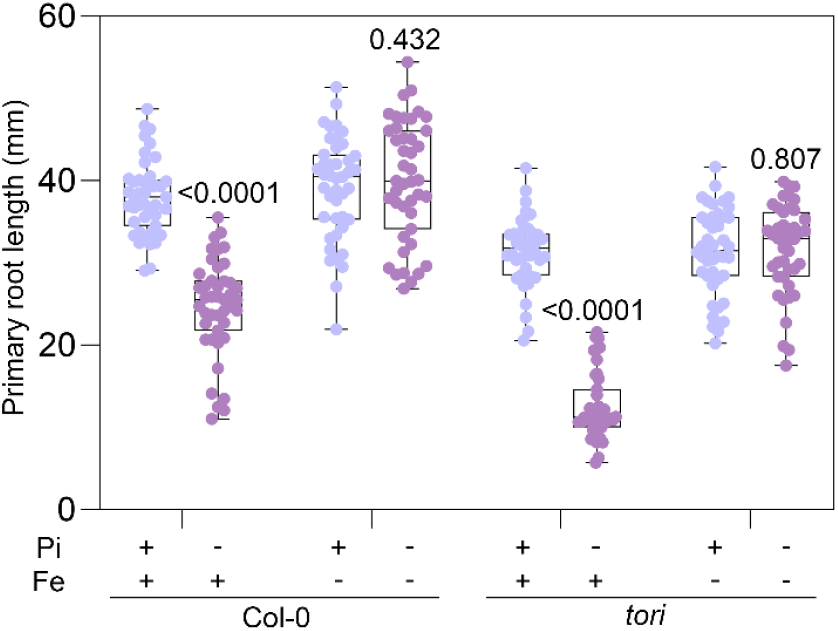
Role of iron in modulating phenotype of phosphate limitation. Primary root length in Col-0 and *tori* growing in high (+, 625 μM Pi) and low (-, 10 μM Pi) P regimes with and without 50 μM Fe. Experiments were repeated three times, and the graphs indicate a biological replicate (n=43, two-way ANOVA with Tukey’s multiple comparisons test, p-values are indicated in the graphs).

**Figure S8.**
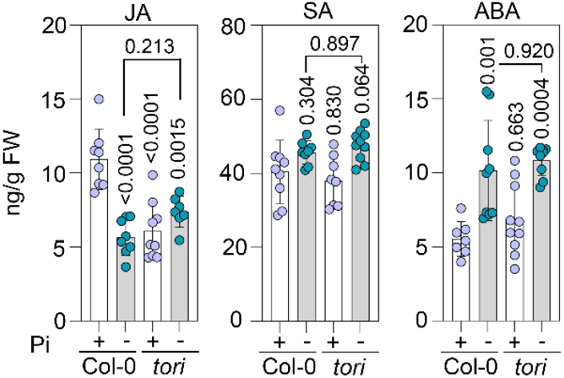
Hormone quantification in different phosphate availability. Estimation of JA, SA and ABA level in Col-0 and *tori* in high and low P regimes (n=7-10, One-way ANOVA with Tukey’s multiple comparisons test, p-values are indicated in the graphs).

## References

1. Castrillo, G., Teixeira, P.J.P.L., Paredes, S.H., Law, T.F., de Lorenzo, L., Feltcher, M.E., Finkel, O.M., Breakfield, N.W., Mieczkowski, P., Jones, C.D., et al. (2017). Root microbiota drive direct integration of phosphate stress and immunity. Nature 2017 543:7646 543, 513–518. 10.1038/nature21417.

2. Paries, M., and Gutjahr, C. (2023). The good, the bad, and the phosphate: regulation of beneficial and detrimental plant–microbe interactions by the plant phosphate status. New Phytologist 239, 29–46. 10.1111/NPH.18933.

3. Rubio, V., Linhares, F., Solano, R., Martín, A.C., Iglesias, J., Leyva, A., and Paz-Ares, J. (2001). A conserved MYB transcription factor involved in phosphate starvation signaling both in vascular plants and in unicellular algae. Genes Dev 15, 2122–2133. 10.1101/gad.204401.

4. Bustos, R., Castrillo, G., Linhares, F., Puga, M.I., Rubio, V., Pérez-Pérez, J., Solano, R., Leyva, A., and Paz-Ares, J. (2010). A central regulatory system largely controls transcriptional activation and repression responses to phosphate starvation in arabidopsis. PLoS Genet 6. 10.1371/journal.pgen.1001102.

5. Jamsheer, M.K., Awasthi, P., and Laxmi, A. (2022). The social network of target of rapamycin complex 1 in plants. J Exp Bot 73, 7026–7040. 10.1093/JXB/ERAC278.

6. De Vleesschauwer, D., Filipe, O., Hoffman, G., Seifi, H.S., Haeck, A., Canlas, P., Van Bockhaven, J., De Waele, E., Demeestere, K., Ronald, P., et al. (2018). Target of rapamycin signaling orchestrates growth-defense trade-offs in plants. New Phytologist 217, 305– 319. 10.1111/nph.14785.

7. Margalha, L., Confraria, A., and Baena-González, E. (2019). SnRK1 and TOR: modulating growth–defense trade-offs in plant stress responses. J Exp Bot 70, 2261–2274. 10.1093/jxb/erz066.

8. Xiong, Y., McCormack, M., Li, L., Hall, Q., Xiang, C., and Sheen, J. (2013). Glucose–TOR signalling reprograms the transcriptome and activates meristems. Nature 496, 181–186. 10.1038/nature12030.

9. Ye, R., Wang, M., Du, H., Chhajed, S., Koh, J., Liu, K. hsiang, Shin, J., Wu, Y., Shi, L., Xu, L., et al. (2022). Glucose-driven TOR–FIE–PRC2 signalling controls plant development. Nature 2022 609:7929 609, 986–993. 10.1038/s41586-022-05171-5.

10. Dobrenel, T., Mancera-Martínez, E., Forzani, C., Azzopardi, M., Davanture, M., Moreau, M., Schepetilnikov, M., Chicher, J., Langella, O., Zivy, M., et al. (2016). The Arabidopsis TOR Kinase Specifically Regulates the Expression of Nuclear Genes Coding for Plastidic Ribosomal Proteins and the Phosphorylation of the Cytosolic Ribosomal Protein S6. Front Plant Sci 7, 1611. 10.3389/fpls.2016.01611.

11. Misson, J., Raghothama, K.G., Jain, A., Jouhet, J., Block, M.A., Bligny, R., Ortet, P., Creff, A., Somerville, S., Rolland, N., et al. (2005). Misson J, Raghothama KG, Jain A, et al. 2005. A genome-wide transcriptional analysis using Arabidopsis thaliana Affymetrix gene chips determined plant responses to phosphate deprivation. Proceedings of the National Academy of Sciences of the United States. Proc Natl Acad Sci U S A 102, 11934–11939. 10.1073/pnas.0505266102.

12. Park, S.H., Jeong, J.S., Huang, C.H., Park, B.S., and Chua, N.H. (2023). Inositol polyphosphates-regulated polyubiquitination of PHR1 by NLA E3 ligase during phosphate starvation response in Arabidopsis. New Phytologist 237, 1215–1228. 10.1111/NPH.18621.

13. Gutiérrez-Alanís, D., Yong-Villalobos, L., Jiménez-Sandoval, P., Alatorre-Cobos, F., Oropeza-Aburto, A., Mora-Macías, J., Sánchez-Rodríguez, F., Cruz-Ramírez, A., and Herrera-Estrella, L. (2017). Phosphate Starvation-Dependent Iron Mobilization Induces CLE14 Expression to Trigger Root Meristem Differentiation through CLV2/PEPR2 Signaling. Dev Cell 41, 555-570.e3. 10.1016/J.DEVCEL.2017.05.009.

14. Qiang, X., Weiss, M., Kogel, K.H., and Schäfer, P. (2012). Piriformospora indica—a mutualistic basidiomycete with an exceptionally large plant host range. Mol Plant Pathol 13, 508–518. 10.1111/J.1364-3703.2011.00764.X.

15. Jogawat, A., Meena, M.K., Kundu, A., Varma, M., and Vadassery, J. (2020). Calcium channel CNGC19 mediates basal defense signaling to regulate colonization by Piriformospora indica in Arabidopsis roots. J Exp Bot 71, 2752–2768. 10.1093/JXB/ERAA028.

16. Trejo-Fregoso, R., Rodríguez, I., Ávila, A., Juárez-Díaz, J.A., Rodríguez-Sotres, R., Martínez-Barajas, E., and Coello, P. (2022). Phosphorylation of S11 in PHR1 negatively controls its transcriptional activity. Physiol Plant 174, e13831. 10.1111/PPL.13831.

17. Miura, K., Rus, A., Sharkhuu, A., Yokoi, S., Karthikeyan, A.S., Raghothama, K.G., Baek, D., Koo, Y.D., Jin, J.B., Bressan, R.A., et al. (2005). The Arabidopsis SUMO E3 ligase SIZ1 controls phosphate deficiency responses. Proc Natl Acad Sci U S A 102, 7760–7765. 10.1073/PNAS.0500778102/SUPPL_FILE/00778SUPPTEXT.PDF.

18. Navarro, C., Mateo-Elizalde, C., Mohan, T.C., Sánchez-Bermejo, E., Urrutia, O., Fernández-Muñiz, M.N., García-Mina, J.M., Muñoz, R., Paz-Ares, J., Castrillo, G., et al. (2021). Arsenite provides a selective signal that coordinates arsenate uptake and detoxification through the regulation of PHR1 stability in Arabidopsis. Mol Plant 14, 1489– 1507. 10.1016/J.MOLP.2021.05.020.

19. Cho, H., Banf, M., Shahzad, Z., Van Leene, J., Bossi, F., Ruffel, S., Bouain, N., Cao, P., Krouk, G., De Jaeger, G., et al. (2023). ARSK1 activates TORC1 signaling to adjust growth to phosphate availability in Arabidopsis. Current Biology 33, 1778-1786.e5. 10.1016/J.CUB.2023.03.005.

20. Dong, Y., Aref, R., Forieri, I., Schiel, D., Leemhuis, W., Meyer, C., Hell, R., and Wirtz, M. (2022). The plant TOR kinase tunes autophagy and meristem activity for nutrient stress-induced developmental plasticity. Plant Cell 34, 3814–3829. 10.1093/PLCELL/KOAC201.

21. Zhou, X., Clister, T.L., Lowry, P.R., Seldin, M.M., Wong, G.W., and Zhang, J. (2015). Dynamic Visualization of mTORC1 Activity in Living Cells. Cell Rep 10, 1767–1777. 10.1016/j.celrep.2015.02.031.

22. Hatakeyama, R., Péli-Gulli, M.P., Hu, Z., Jaquenoud, M., Garcia Osuna, G.M., Sardu, A., Dengjel, J., and De Virgilio, C. (2019). Spatially Distinct Pools of TORC1 Balance Protein Homeostasis. Mol Cell 73, 325-338.e8. 10.1016/j.molcel.2018.10.040.

23. Xiong, Y., and Sheen, J. (2012). Rapamycin and glucose-target of rapamycin (TOR) protein signaling in plants. J Biol Chem 287, 2836–2842. 10.1074/jbc.M111.300749.

24. Deprost, D., Yao, L., Sormani, R., Moreau, M., Leterreux, G., Nicolaï, M., Bedu, M., Robaglia, C., and Meyer, C. (2007). The Arabidopsis TOR kinase links plant growth, yield, stress resistance and mRNA translation. EMBO Rep 8, 864–870. 10.1038/sj.embor.7401043.

25. Karthikeyan, A.S., Varadarajan, D.K., Jain, A., Held, M.A., Carpita, N.C., and Raghothama, K.G. (2007). Phosphate starvation responses are mediated by sugar signaling in Arabidopsis. Planta 225, 907–918. 10.1007/S00425-006-0408-8/FIGURES/8.

26. Ames, B.N. (1966). [10] Assay of inorganic phosphate, total phosphate and phosphatases. Methods Enzymol 8. 10.1016/0076-6879(66)08014-5.

27. Livak, K.J., and Schmittgen, T.D. (2001). Analysis of relative gene expression data using real-time quantitative PCR and the 2-ΔΔCT method. Methods 25, 402–408. 10.1006/meth.2001.1262.

28. Vadassery, J., Reichelt, M., Hause, B., Gershenzon, J., Boland, W., and Mithöfer, A. (2012). CML42-Mediated Calcium Signaling Coordinates Responses to Spodoptera Herbivory and Abiotic Stresses in Arabidopsis. Plant Physiol 159, 1159. 10.1104/PP.112.198150.

29. Clarke, J.D. (2009). Cetyltrimethyl Ammonium Bromide (CTAB) DNA Miniprep for Plant DNA Isolation. Cold Spring Harb Protoc 2009, pdb.prot5177. 10.1101/pdb.prot5177.

